# From behaviour to complex communities: Resilience to anthropogenic noise in a fish-induced trophic cascade

**DOI:** 10.1101/2022.07.05.498792

**Authors:** Emilie Rojas, Mélanie Gouret, Simon Agostini, Sarah Fiorini, Paulo Fonseca, Gérard Lacroix, Vincent Médoc

## Abstract

Sound emissions from human activities represent a pervasive environmental stressor. Individual responses in terms of behaviour, physiology or anatomy are well documented but whether they propagate through nested ecological interactions to alter complex communities needs to be better understood. This is even more relevant for freshwater ecosystems that harbour a disproportionate fraction of biodiversity but receive less attention than marine and terrestrial systems. We conducted a mesocosm investigation to study the effect of chronic exposure to motorboat noise on the dynamics of a freshwater community including phytoplankton, zooplankton, and roach as a planktivorous fish. As expected under the trophic cascade hypothesis, roach predation induced structural changes in the planktonic communities. Surprisingly, although roach changed their feeding behaviour in response to noise, the dynamics of the roach-dominated planktonic communities did not differ between noisy and noiseless mesocosms. This suggests that the top-down structuring influence of roach on planktonic communities might be resilient to noise and reveals the difficulties on extrapolating impacts form individual responses to complex communities.

## 1. INTRODUCTION

The trophic cascade, one of the most influential concepts in ecology, specifies the effects of predators that propagate downward through food webs across multiple trophic levels [1,2]. Considering a series of nested consumer-resource interactions (i.e., a food chain), top predators have a direct negative effect on mesopredators and indirect positive and negative effects alternatively on lower trophic levels. Top-down cascade effects can result from changes in predator density (density-mediated trophic cascade) or behaviour (trait-mediated trophic cascade) and much attention has focused on identifying the intrinsic and extrinsic determinants of their strength [3,4]. In particular, this helped to better understand the structural impact of several anthropogenic stressors including warming, salinization, chemical pollution or habitat degradation [5–7].

Noise emissions from transportation, cities, industry, military and recreational activities represent another pervasive anthropogenic stressor [8,9]. They span all ecosystems even in the most remote places [10] and have been shown to alter communication, social interactions, use of space, activity patterns, foraging and reproduction in a wide range of taxa [11–13]. First evidence of the cascading effects of noise pollution came from the long-term investigations conducted in the natural gas fields of northwest New Mexico. Bird response to gas-well-compressor noise was found species-specific with some key seed dispersers (mountain bluebirds and Woodhouse’s scrub-jays) avoiding noisy areas where pollinators like hummingbirds had on the contrary higher reproductive success [14]. Long-term consequences include alterations in plant communities that persist after removal of the noise source [15]. The propensity of anthropogenic noise to indirectly affect species, and typically primary producers, through a series of nested direct interactions had also been suggested experimentally. Barton *et al*.[16] exposed a three-level terrestrial food chain to various soundscapes for 14 days in plant growth chambers and found that urban sounds and rock music made lady beetles less effective predators, reducing the strength of top-down control on aphids, whose density increased. More aphids ultimately resulted in reduced soybean biomass. Although freshwaters harbour a disproportionate fraction of earth’s biodiversity [17] and suffer a greater decline in species richness compared to terrestrial and marine habitats [18,19], they often receive less attention, and research on the impacts of noise pollution is no exception. For instance, we known that fish responses to noise include changes in behaviour and abundances [20–23] but whether these effects spread along food webs to alter planktonic communities through cascading effects remains to be investigated. Similarly, the response of freshwater plankton to noise is largely overlooked. Available evidence to date comes from water fleas, *Daphnia* spp., which are widespread pelagic crustaceans (Cladocera) and an important source of food for upper trophic levels [24,25]. Surprisingly, knowing that marine invertebrates of similar size were found to adjust their swimming activity in response to natural or artificial sounds [26], water fleas exposed to band-pass filtered white noise either continuous or intermittent did not show any alteration in swimming speed or depth [27,28]. However, long-term effects of chronic exposure on their survival and reproductive success have yet to be explored, and overall, we lack knowledge on the dynamics of plankton under anthropogenic noise. Here we conducted a mesocosm investigation to study the temporal dynamics of a zooplankton – phytoplankton system under the presence or absence of a planktivorous fish, with and without exposure to motorboat noise. We used the roach *Rutilus rutilus* as top predator: a widespread Eurasian cyprinid fish whose response to motorboat noise has been documented. Roach have specialized hearing structures, the Weberian ossicles, that conduct sound from the swim-bladder to the inner ear and provide high sensitivity to sound pressure [29–31]. They can detect sounds between 10 Hz and 5 kHz with a maximum sensitivity of 60 dB re. 1 μPa between 500 and 1000 Hz [32]. Roach have been found to respond to authentical motorboat sounds with fewer feeding attempts, higher latency to enter the open area and longer time spent in the vegetation, and these effects persisted after five days of exposure suggesting the absence of habituation [33]. After our mesocosm investigation, and in order to get insights into fish growth and behaviour, the roach have been collected, weighted, and measured, and moved to aquaria to assess prey consumption, mobility and group cohesion in the presence or absence of boat noise. Given the lack of knowledge on how freshwater plankton respond to chronic noise exposure, we had no clear prediction on the effect of boat noise on the dynamics of the zooplankton – phytoplankton system without fish. Under the trophic cascade hypothesis, we expected roach presence to reduce the top-down control of phytoplankton by zooplankton. Considering that motorboat noise was found to negatively influence foraging in roach [33], we predicted a decrease in the strength of top-down cascading effect. Concerning roach behaviour, we expected behavioural alterations consistent with the weakening of the trophic cascade in the noisy mesocosms.

## 2. MATERIALS AND METHOD

The investigation has been conducted at the PLANAQUA experimental platform of the CEREEP-Ecotron Île-de-France research station (48° 16′10.92 N, 2° 43′50.879 E, Seine et Marne, France) and lasted six weeks (August 31 – October 14, 2020), which corresponds to a prolonged exposure to motorboat noise following Johansson et al. (2016). We assigned three mesocosms to one of four treatments (*N* = 3 replicates, 12 mesocosms in total): (1) no fish - no noise, (2) no fish - noise, (3) fish – no noise, and (4) fish – noise.

### 2.1 Preparation of the mesocosms and animal collection

Each mesocosm (3.4 m diameter, 1.1 m depth) was filled with 9,079 L of water from the storage lakes of the field station that naturally host zooplankton and phytoplankton communities. An underwater loudspeaker (Electrovoice UW30) was submerged five cm below the surface in the center of each mesocosm. It was connected to an amplifier (Dynavox CS-PA 1MK) and then to an audio player (Handy’s H4n zoom), both placed inside a waterproof electric box next to the mesocosm. To promote the growth of phytoplankton, we added 30 mL of Algoflash^®^ (41.4 μg of phosphorus, nitrogen, and potassium per liter) at the beginning of the investigation (August 20) and 50 additional mL at the middle of the investigation (September 19) after we detected a drop in the amount of chlorophyll in the mesocosms using a multiparameter probe (YSI ExO-2). From March to April 2020, we used fish traps to gradually collect roach from the storage lakes of the PLANAQUA platform, and we stored them in a pond containing the same planktonic communities as the mesocosms. These fish were the descendants of roach used in previous investigations conducted on the platform. They grew in quiet conditions and had never experienced motorboat noise before. At Day 0, 96 roach of similar size (8.54 ± 2.32 cm for standard length, SL) were randomly collected from the storage pond using a seine net, measured, and weighted to the nearest 0.01 cm and 1 g, and distributed in groups of 16 between the six mesocosms (fish – no noise and fish – noise treatments) so as to homogenize size distribution and total biomass between the mesocosms. We placed anti-bird nets on top of the mesocosms to avoid avian predation.

### 2.2 Plankton dynamics

To assess plankton dynamics, we sampled the mesocosms 13 times from Day 0 to Day 42 and every two or four days. The temporal variation of phytoplankton was assessed through the quantification of green algae, cyanobacteria, and diatom densities. We sampled eight liters of water per mesocosm using a 2-L sampling bottle (Uwitec) at four different positions. Analyses were made in the laboratory using a BBE FluoroProbe™ spectrofluorometer (BBE Moldaenke GmbH, Schwentinental) on a 125-mL subsample previously kept in the dark for one hour. For detecting potential top-down effects on zooplankton, we choose to focus on mesoplankton organisms (Cladocera, copepodits and adults of Copepods, and *Chaoborus* larvae), which appeared as the most responsive organisms in previous mesocosm experiments realized in comparable conditions [34]. To assess their temporal variation in zooplankton, we sampled 24 liters of water using a 2-L sampling bottle at twelve different positions and depths in each mesocosm. Water was filtered with a 50-μm nylon filter and zooplankton fixed in 15-mL of 90% ethanol. Taxa identification and counting was made on a 3-mL subsample following [35] for cladocerans and copepods (nauplii not counted), and [36] for aquatic insects.

### 2.3 Fish growth and behaviour

The behavioural tests took place in an experimental room of the PLANAQUA platform thermo-regulated at 17°C. We equipped four 110-L aquaria (80 cm length × 35 cm width × 40 cm height) with an underwater loudspeaker (Electrovoice UW30) in the middle of the left end surrounded by acoustic foam (1.5-cm thick) to attenuate vibrations, a neon light, and a camera (HD-TVI ABUS TVVR33418) above, and black plastic boards outside to avoid visual contacts with the experimenters that may provide stress. The speaker was connected to an amplifier (Dynavox CS-PA 1MK) and to an audio player (Handy’s H4n zoom). The aquaria were filled with water from the control mesocosms (no fish – no noise treatment) filtered through a 50-μm nylon mesh filter to remove zooplankton. This experimental design allowed us to run four tests simultaneously with one treatment *per* aquarium depending on the noise condition in the mesocosm and later in the aquarium (“mesocosm – aquarium” noise conditions): (1) no noise – no noise, (2) no noise – noise, (3) noise – no noise and (4). Between two consecutive runs of tests, each aquarium was assigned another treatment to avoid an effect of the aquarium, while water was changed to remove chemical cues.

At the end of the mesocosm investigation (Day 44), roach were removed from the mesocosms using a seine net and measured and weighted to the nearest 0.01 cm and 1 g. We matched these values with the weights and lengths measured at the beginning of the mesocosm investigation to recognize fish and calculate individual growth. Roach were then randomly assigned to groups of three individuals and moved into one of the four experimental aquaria. In total, we formed 28 groups (*N* = 7 replicates per treatment) with two or three groups per mesocosm. Once in the aquarium, they first experienced ambient noise during an acclimatization period of one hour, and then 40 minutes of either ambient noise or ambient noise supplemented with motorboat sounds, depending on the treatment (see section 2.4 for further detail on the playback tracks). At the middle of the exposure period (i.e. after 20 min), we introduced 50 *Chaoborus* larvae (Diptera) and 50 *Daphnia* sp. (Crustacera: Cladocera), previously collected from the control mesocosms using a 2-L sampling bottle and 50-μm nylon mesh filter. Both invertebrates are common prey of roach, differ in terms of size and mobility, and were found in the mesocosms. Although *Chaoborus* larvae are natural predators of *Daphnia* [37], we expected no predation events considering the short duration of the experiment. At the end of the experiment, roach and invertebrates were removed from the aquarium, counted, and returned to the storage pond. The videos were analyzed using Kinovea v. 0.9.4 to get the xy coordinates of each fish each second and during each boat sound (for a total of 691s and corresponding to ambient noise for the other treatment), and then the following parameters were calculated: (1) the cumulative swimming distance (total distance covered by the three individuals) as a proxy of mobility, (2) the distance between the barycenter of each group and the center of the speaker as a proxy of aversion to noise, and (3) the area occupied by the group as a proxy of group cohesion.

Group’s barycenter was calculated using individual coordinates as follows:

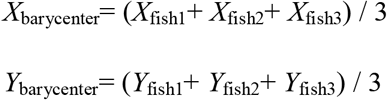

The distance between the barycenter and the loudspeaker (*D* in cm) was calculated as follows:

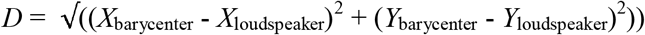

Group’s area (*A* in cm^2^) was calculated as follows:

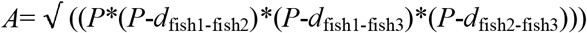

Where *d* corresponds to the distance between two fish in cm and *P* to the perimeter of the group in cm with:

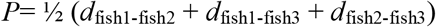

Group barycenter and area were calculated every second during the boat sounds and then averaged.

The behavioural tests took place from 8 am to 6 pm and needed two consecutive days with four mesocosms (two from the fish – no noise treatment and two from the fish – noise treatment) processed on Day 43 and the two others (one *per* treatment) on Day 44. We also conducted four additional tests (two *per* noise condition) without roach to control for *Chaoborus* predation on *Daphnia* and overall invertebrate mortality in the absence of predation.

### 2.4 Noise treatments

We used Audacity 2.2.1 to generate the audio tracks and an Aquarian Audio H2A-HLR hydrophone (frequency response from 10 to 100 kHz) connected to a ZOOM H4next Handy recorder for all the recordings. The level of background noise did not differ between the mesocosms and ranged from 90 to 95 dB re. μPa. In the mesocosms without boat noise, a 1-hr audio track of silence was looped continuously. In the mesocosms with boat noise, we used the audio tracks described in [38] where 150 sounds of small recreational boats have been distributed over the nine consecutive 1-hr audio tracks of silence going from 9 a.m. to 6 p.m. so as to mimic the daily activity of a small leisure base (Fig. 1A and Supp. Mat. 1, see Rojas et al. 2021 for more details on the audio tracks and the original recordings). We broadcasted silence the rest of the time. We applied a linear fading on both ends of the boat sounds to make them emerge from background noise and adjusted their levels with Audacity to obtain naturally occurring signal-to-noise ratios (SNR) ranging from 4.81 to 27 dB. We used the SNR function of the *seewave* R package [39]:

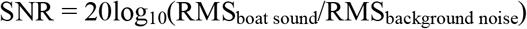

where RMS is the root-mean-square sound pressure of either the re-recordings of the boat sounds in the mesocosm or the recording of background noise.

**Figure 1:**
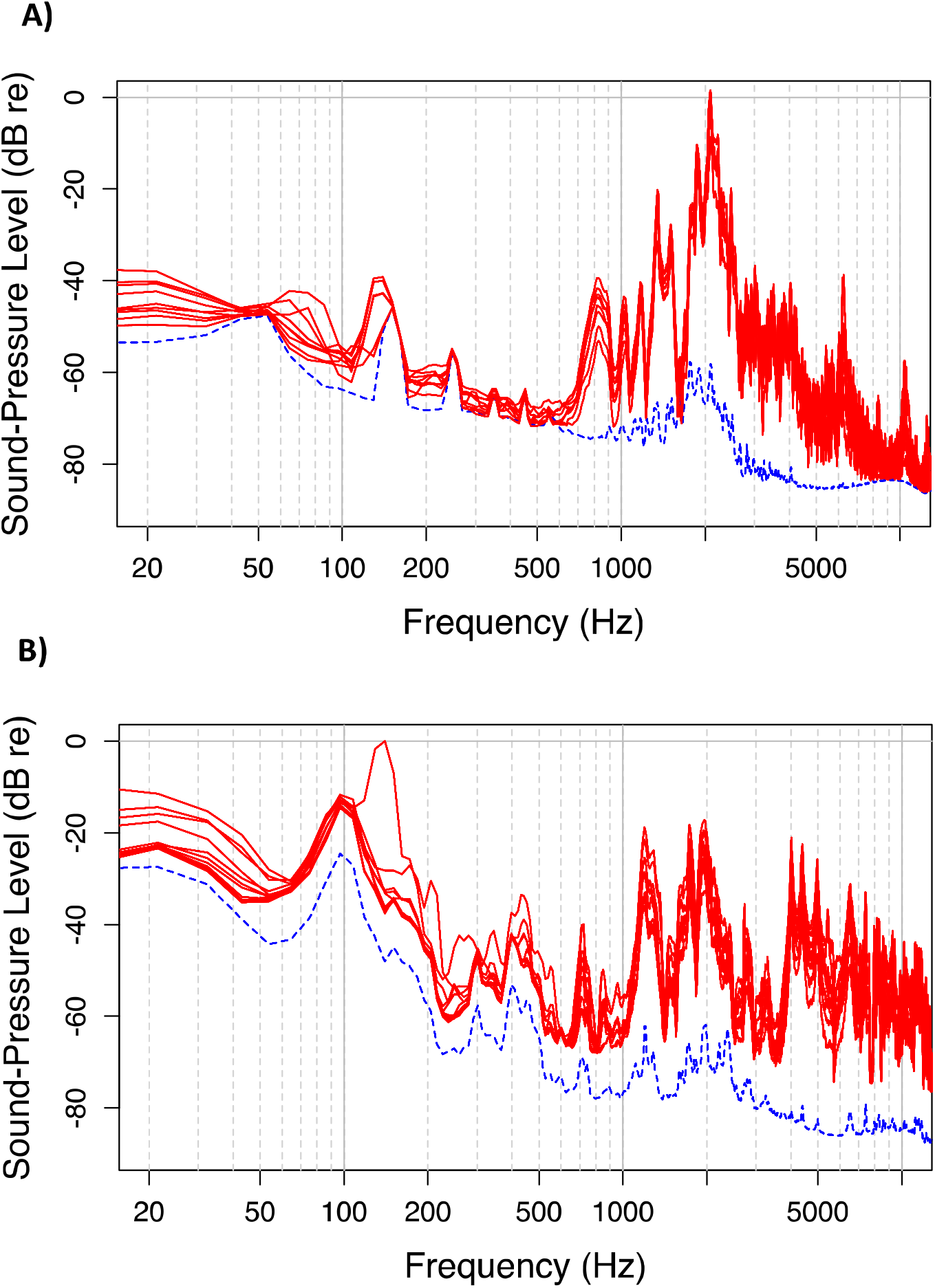
Sound spectra of the two noise conditions (control: dashed lines, boat noise: one solid lines for each hour from 9 am to 6 pm) broadcasted in A) the mesocosms and B) the aquaria. Spectra were made from 1-hour recordings in the mesocosms and 20-min recordings in the aquaria.

In the aquaria without boat noise and to encompass the 1-h acclimatization period and the 40-min exposure period, we used a 100-min audio track of background noise previously recorded in the center of one mesocosm from the no noise – no fish treatment. We adjusted sound level to match that of the mesocosms. In the aquaria with boat noise, we used the 100-min audio track of background noise to which we added twelve boat sounds randomly selected from those broadcasted in the mesocosms. We randomly distributed the sounds over the 40-min exposure period and adjusted their level to match the range of SNR values we had in the mesocosms (approx. 4.81 to 27 dB). In terms of boat traffic, this acoustic regime was representative of the highest activity that roach experienced within a day in the mesocosms (Fig. 1B and Supp. Mat. 1).

### 2.5 Data analysis

All the statistics were performed using R [40] with a significance level of 0.05. We used the Principal Response Curve (PRC, *prc* function of the *vegan* R package [41]) to study how the planktonic communities exposed to roach, boat noise, or both, have diverged over time compared to control communities (i.e. from the no fish – no noise treatment). PRC is a special case of redundancy analysis including time-series data particularly suited to the study of community dynamics in mesocosms [42]. It typically results in a diagram with one curve for each treatment, the time on the x-axis, the first major component of the community effects on the left y-axis and the weights of the taxa on the right y-axis. The more the weight deviates from zero the more the corresponding taxon contributes to the deviation from the control. We used Hellinger-transformed taxa (square root of the relative abundance) to reduce the influence of both rare (low abundances and/or many zeros) and abundant taxa [43]. Significance in the PRC was tested using a permutation test (*anova.cca* function of the *vegan* R package) accounting for the non-independence of data due to repeated measurements on the same mesocosm. Significance in the difference between each treatment and the control was assessed with a multiple comparison test (*multiconstrained* function of the *BiodiversityR* R package).

Roach’ growth rate (*G*) was computed using the formulae:

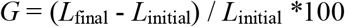

where *L*_final_ and *L*_initial_ are the final and initial SL of roach, respectively.

Because growth data met the normality and homoscedasticity assumptions (Shapiro-Wilk and Bartlett tests, all *p* values > 0.05), we used a linear mixed-effect model (*lme4* R package [44]) with the noise condition as fixed factor and the mesocosm as random factor to test for significance of the difference in growth rate between the treatments.

The effects of noise and pre-exposure to noise on roach predation were assessed in two ways. First, we used a generalized linear mixed-effect model assuming a Poisson distribution to explain the total number of prey eaten as a function of the noise condition in the mesocosm, the noise condition in the aquarium and their interaction as fixed factors, and the mesocosm ID as random factor. Second, we estimated the preference of roach for *Daphnia* over *Chaoborus* larvae using the Manly’s alpha (*α*) preference index [45,46]:

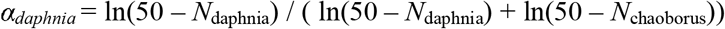

where 50 is the initial number of each prey, and *N*_daphnia_ and *N*_chaoborus_ the numbers of daphnia and *Chaoborus* larvae eaten. The Manly’s alpha accounts for prey depletion during the predation test and ranges from zero when only the alternative prey (here *Chaoborus* larvae) is eaten to one when only the focal prey (here *Daphnia*) is eaten. The value of 0.5 indicates a lack of preference. As recommended by [47], we compared obtained values to the theoretical value of 0.5 using *t* tests except for the “no fish – noise” treatment where we used a Wilcoxon test to deal with the non-normality of data.

Roach behaviour was analyzed with model averaging and an information-theory approach. We used linear mixed-effect models to model each of the three response variables (cumulative swimming distance, distance to the speaker and area of the group) as a function of the noise condition in the mesocosm, the noise condition in the aquarium, the time, and taking the mesocosm ID as random factor as several groups came from the same mesocosm. Because the noise condition that roach have experienced in the mesocosm might have changed their response to boat noise in the aquarium and because the effect of time may vary with the noise condition, we also included the interactions between the two noise treatments and between each noise treatment and time in the predictors. All predictors were centered and scaled using the *standardization* function of the *arm* R package. We ranked all the submodels based on small sampled-corrected AIC values (AIC_c_, *dredge* and *get.models* functions of the *MuMln* Rpackage [48]) and performed model averaging on a confidence set of models using a cut-off of 10 AIC_c_ (*model.avg* function of the *AICcmodavg* R package). The predictors whose parameter estimate had a 95% confidence interval (CI) that included the value of zero were considered as having no significant effect.

## 3. RESULTS

The pelagic zooplankton communities of the mesocosms included four cladoceran families (Bosminidae, Daphniidae, Sididae and Chydoridae), cyclopoid and calanoid copepods, dipteran larvae of the genus *Chaoborus*, and some ostracods captured in the pelagic zone although being mainly benthic organisms. The diagram of the PRC analysis illustrates how adding roach, boat noise or both make the planktonic communities gradually deviate over time from those of the no fish – no noise treatment (i.e., control) considered as the baseline (Fig. 2). Adding boat noise made the communities deviate from the control and adding roach induced a larger deviation without variation between the two noise conditions (i.e., similar trajectories, Fig. 2). Table 1A shows that 30% of total variance was attributed to time and 34% to the treatment regime, including its interaction with time. On the basis of the permutation tests, the treatment regime as well as time and their interaction had a significant influence on the community dynamics (Table 1B). The pairwise comparisons revealed that the difference between the two treatments with roach (with or without noise) was the only to be not significant (Table 1C). Bosminidae and in a lesser extent Chydoridae, both copepod taxa and green algae are indicated with a positive taxon weight in the PRC, suggesting they were expected to increase in abundance with the treatments relative to the control and in proportion to their weight. On the other hand, Daphniidae and, to a lesser extent ostracod, Sididae and *Chaoborus* larvae exhibited negative species weights and were expected to decrease in abundance with the treatments. The taxa weights of diatoms and cyanobacteria were the smallest and close to zero (Fig. 2).

**Figure 2:**
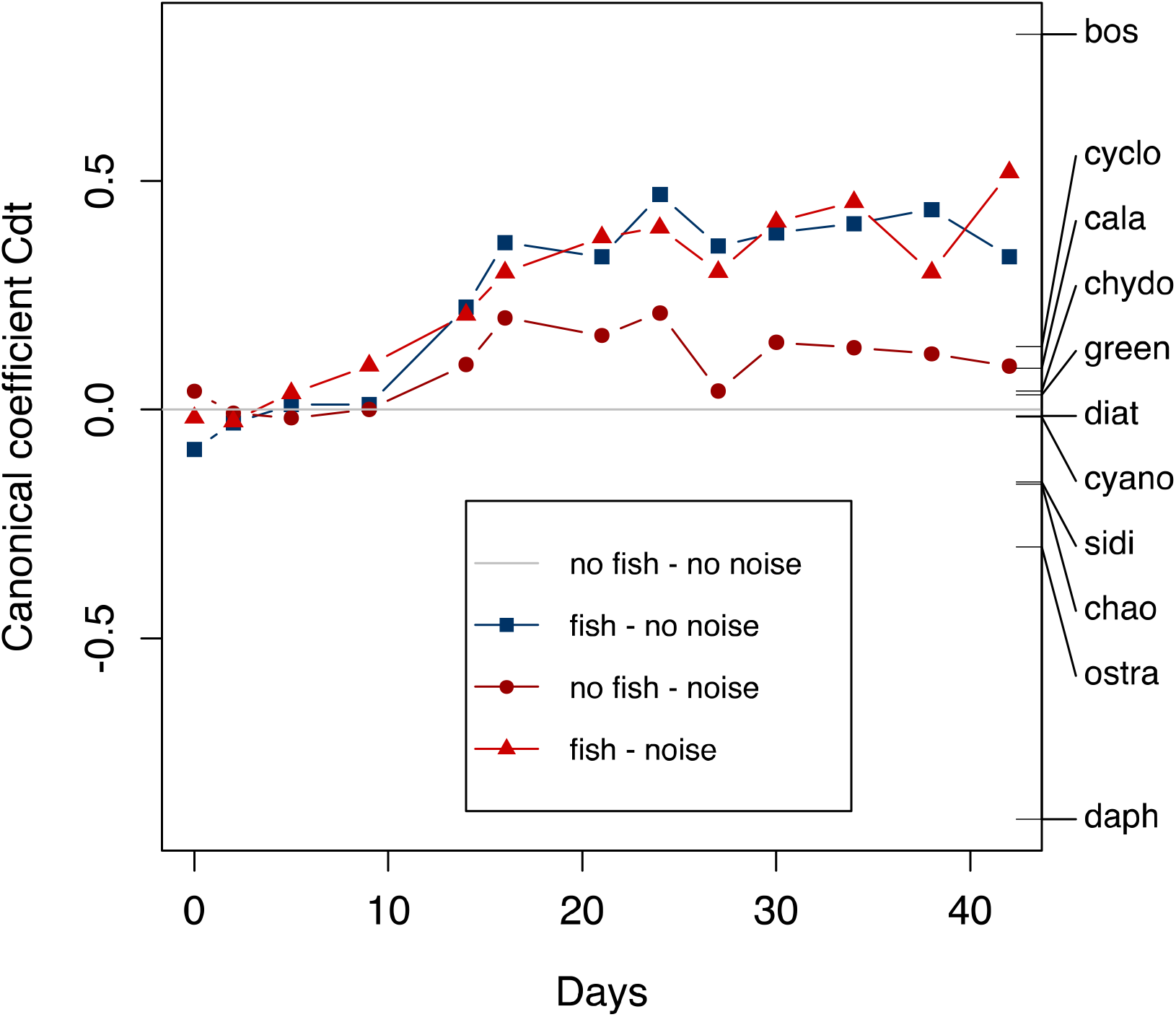
Principal Response Curve (PRC) showing the effects of adding fish (squares), motorboat noise (dots) or both (triangles) on freshwater plankton communities compared to control communities (no fish – no noise, horizontal line, see text for further detail). Species weights are on the left axis (bos: Bosminidae, cyclo: cyclopoid copepods, cala: calanoid copepods, chydo: Chydoridae, green: green algae, diat: diatoms, cyano: cyanobacteria, sidi: Sididae, chao: *Chaoborus* larvae, ostra: ostracods, daph: Daphniidae). See Table 1 for the percentages of variance accounted for and the significance levels.

**Table 1:**
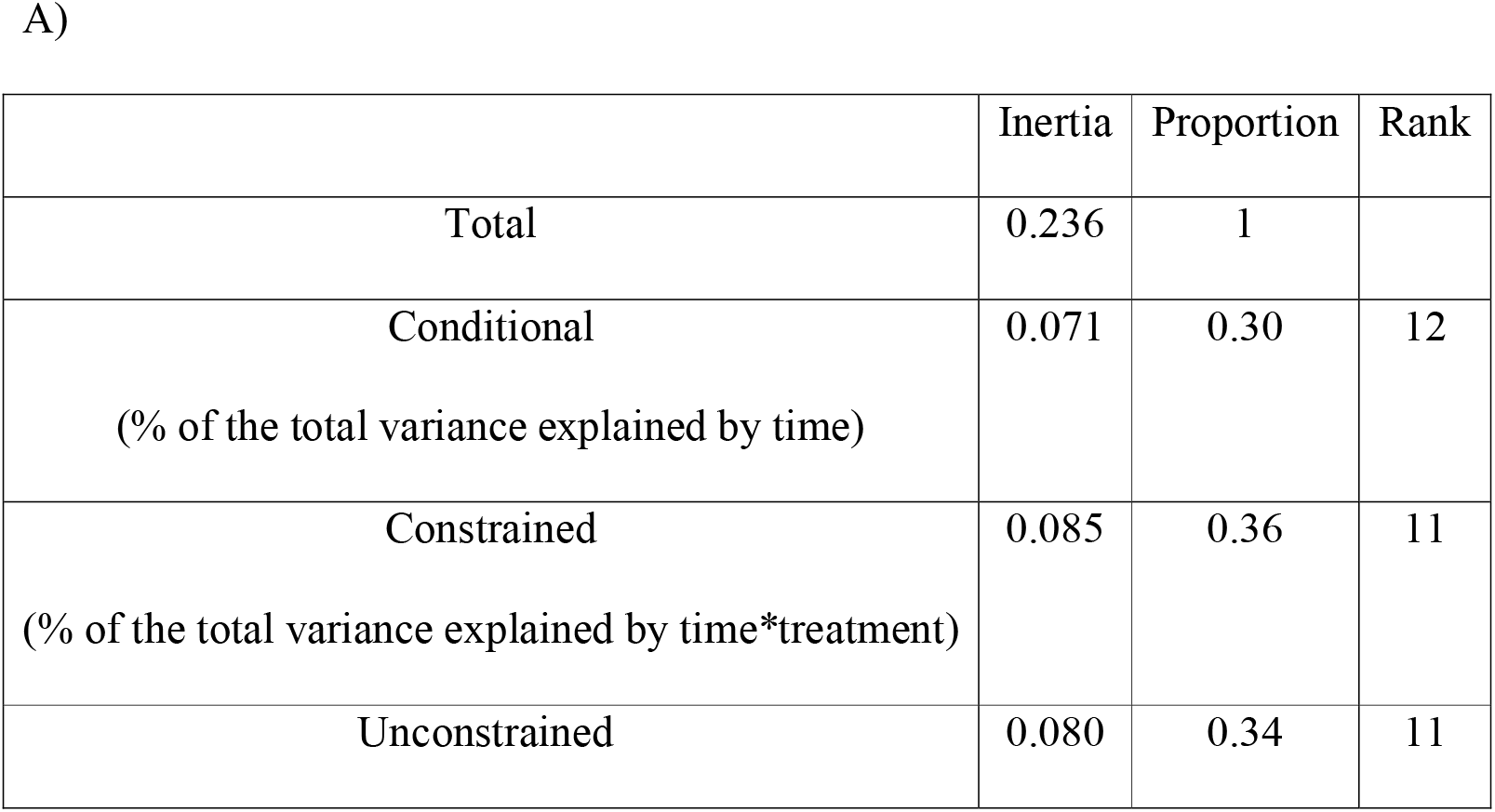
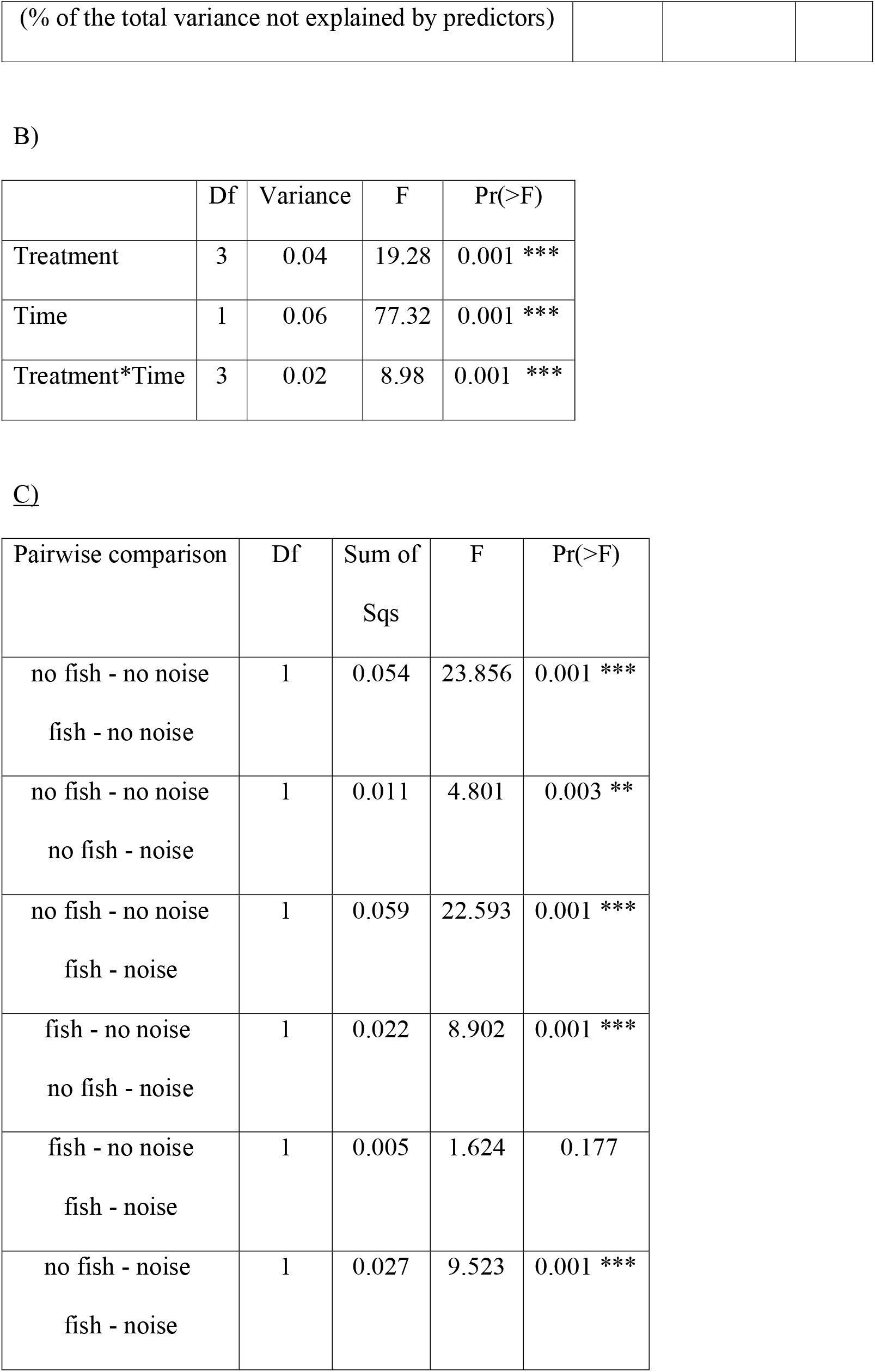
Results of the Principal Response Curve (PRC) for the effect of motorboat noise (absence / presence) and fish (absence / presence, for a total of four treatments) on freshwater planktonic communities. A) Proportion of the total variance explained by the constraints: time, treatment, and their interaction, captured by the canonical 1st axis of the PRC. B) Significance of the PRC diagram on the basis on the permutation test for Constrained Correspondence Analysis (CCA, 999 permutations). C) Pairwise comparisons for all the possible treatment combinations following a CCA analysis.

Individual fish growth did not significantly differ between the noise conditions (*X*^2^_1_ = 1.4813 and *p* = 0.2236, Fig. 3). In the absence of roach, prey survival in the aquaria was 100% for both prey in the absence of boat noise, and 100% for *Chaoborus* larvae and 98% for *Daphnia* in the presence of boat noise. We therefore considered prey mortality during the predation tests to be the result of fish predation only. The effect of noise on the total number of consumed prey depended on the noise condition in the mesocosms, with significantly less prey consumed only for roach coming from the noiseless mesocosms (*X*^2^_1_ = 18.36 and *p* < 0.001 for the interaction between the two noise conditions, Fig. 4A). Whatever the noise condition in the mesocosm, the Manly’s alpha index did not differ from the theoretical value of 0.5 in noiseless aquaria (*t* = 1.36 and *p* = 0.22 for ambient noise, *t* = −0.32 and *p* = 0.76 for boat noise) but was significantly higher with boat noise (*V* = 27 and *p* = 0.03 for ambient noise, *t* = 3.10 and *p* = 0.02 for boat noise, Fig. 4B). Concerning the cumulative swimming distance and the distance to the speaker, the 95% CI of the parameter estimate included the value of zero for all the predictors. Concerning the area occupied by the group, boat noise in the aquarium and time were the only predictors whose parameter estimate 95% CI did not include the value of zero, with significantly positive values (Fig. 5).

**Figure 3:**
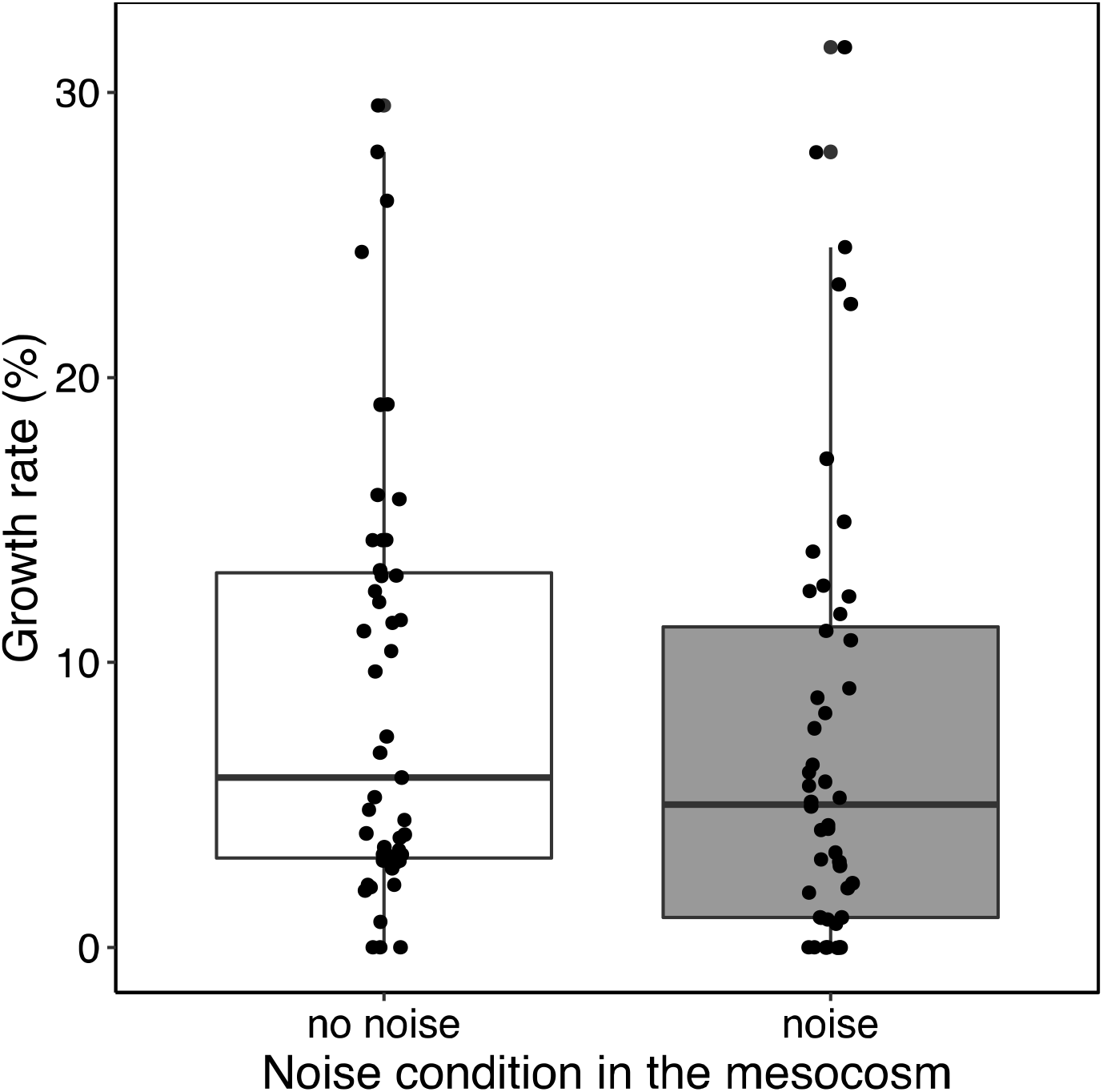
Fish growth rates (medians and interquartile ranges) in the mesocosms depending on the noise condition (white box: ambient, grey box: boat noise, *n* = 48 *per* condition).

**Figure 4:**
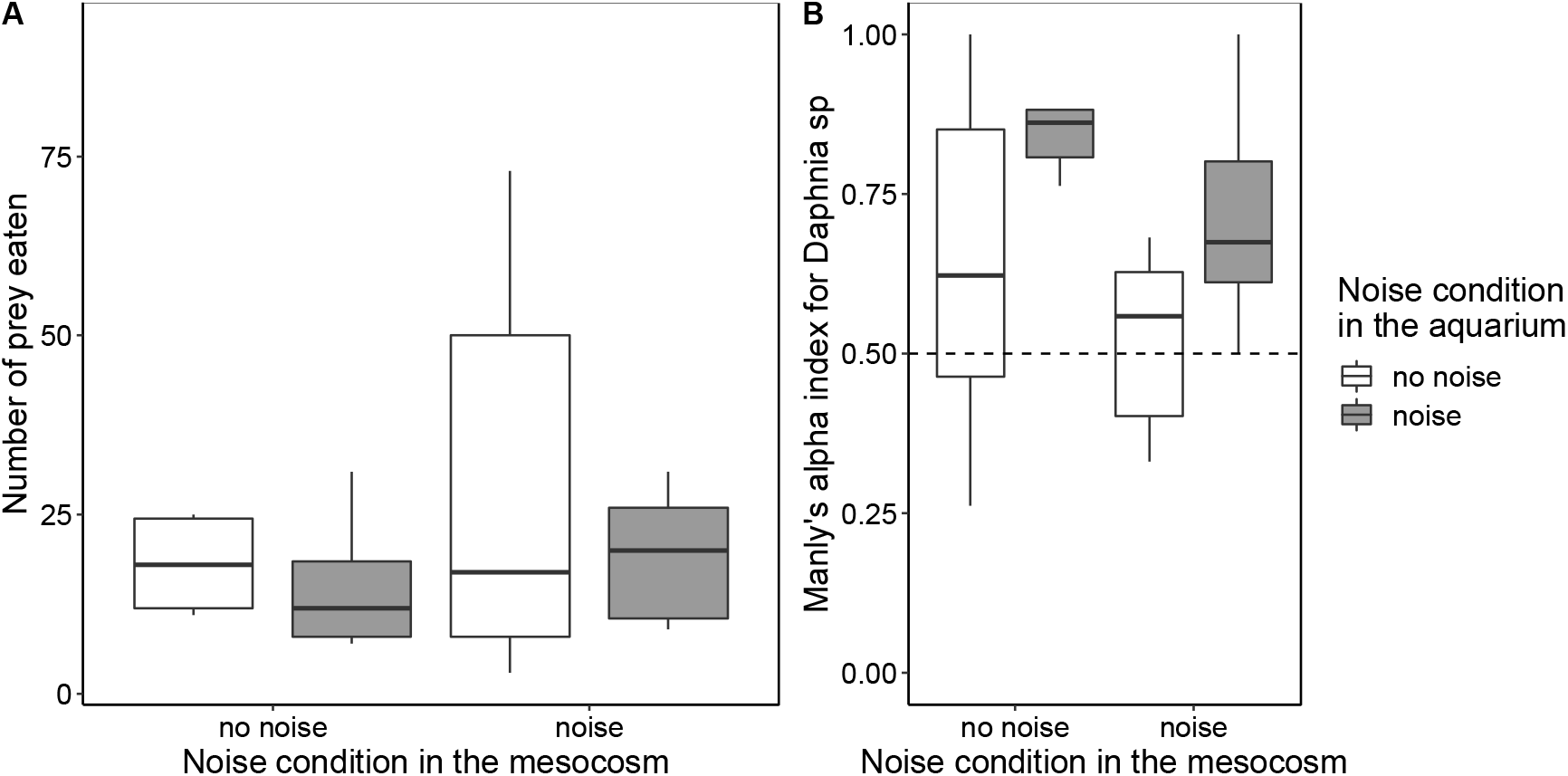
Results of the predation tests in aquaria with A) the total number of prey eaten *per* group of three fish (n=7 groups per acoustic mesocosm-aquarium treatment) presented to 50 *Daphnia* and 50 *Chaoborus* larvae and B) the Manly’s alpha preference index for *Daphnia* over *Chaoborus* larvae (medians and interquartile ranges) as a function of the noise condition previously experienced in the mesocosms and the noise condition in the aquaria (white box: ambient, grey box: boat noise).

**Figure 5:**
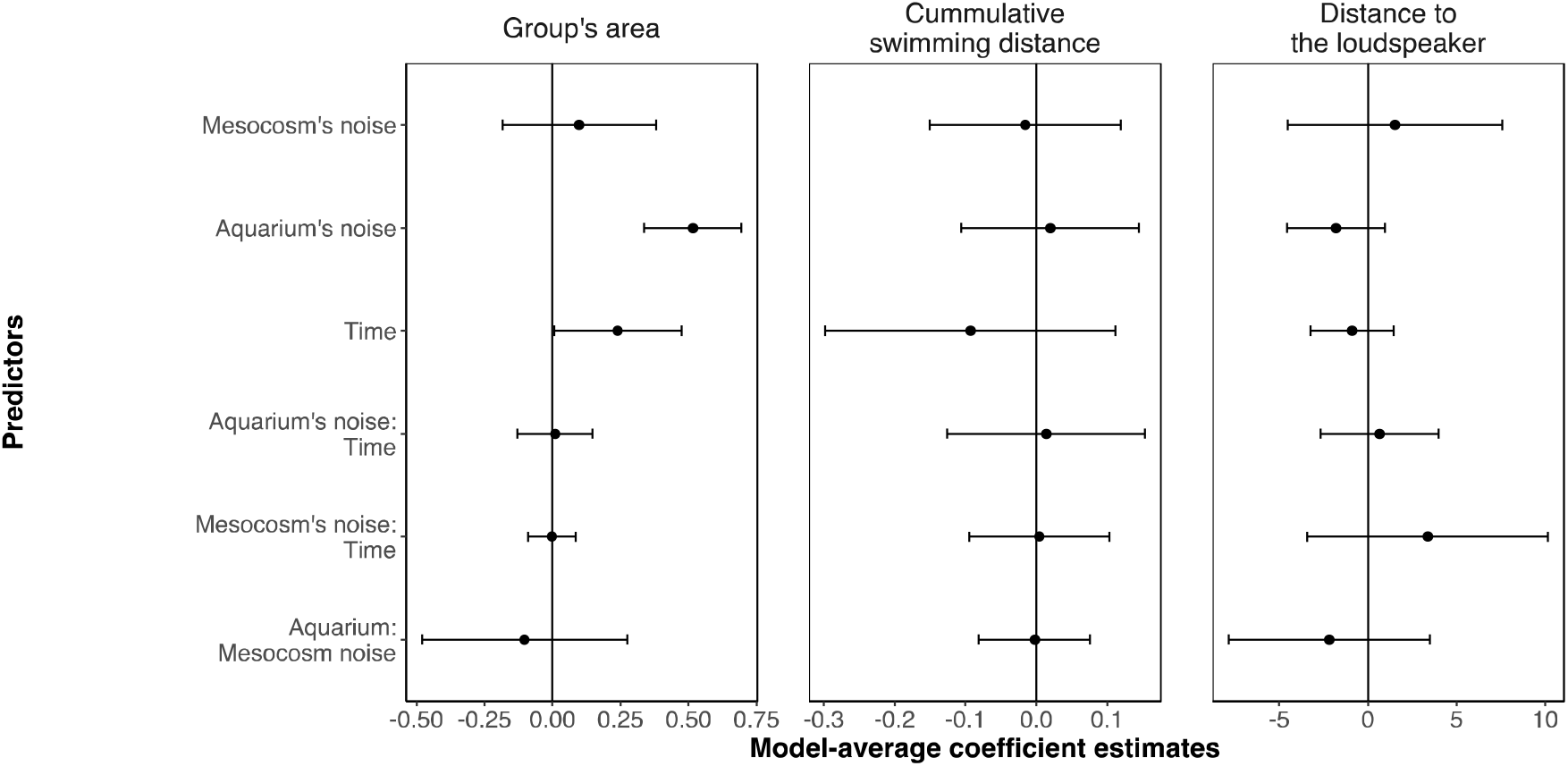
Model-averaged coefficient estimates and 95% confidence intervals for the predictors included in the confidence set of models explaining the behaviour of roach *Rutilus rutilus* in groups of three individuals when feeding on *Daphnia* and *Chaoborus* larvae during the predation tests in aquaria. We used as response variables A) the area of the group, B) the cumulative swimming distance and C) the distance to the speaker. Predictors correspond to the noise condition in the mesocosms (boat noise or ambient noise as control), the noise condition in the aquaria of the predation tests (boat noise or ambient noise as control), the time, and the two-way interactions between time and the noise condition in the two experimental units and between the noise conditions of the two experimental units.

## 4. DISCUSSION

Exposure to anthropogenic noise is known to elicit physiological or behavioural responses in individual organisms [8,11–13,22]. But to what extent these alterations spread across ecological interactions to alter community dynamics and ecosystem functions is not clear. We conducted a mesocosm investigation to study the impact of chronic exposure to motorboat noise on the dynamics of a zooplankton – phytoplankton system either alone or dominated by a planktivorous fish. Although we detected alterations in fish feeding and behaviour, the strength of top-down control and its consequences on the structure of the planktonic communities were resilient to motorboat noise. This suggests that individual responses to noise do not necessarily result in ecological effects at the level of communities.

The pelagic mesoplankton communities of our mesocosms were dominated by cladocerans and copepods, two major groups of herbivorous microcrustaceans widespread in freshwater bodies. In smaller proportions, they also included ostracods (coming from the benthic areas of the mesocosms), and predatory larvae of the *Chaoborus* genus, known to feed on small zooplankton [49]. In fishless mesocosms, daphnid cladocerans gradually became the most abundant taxa. Compared to copepods, cladocerans have higher reproduction rates. Moreover, because daphnids are the largest cladocerans, they suffer smaller predation risk by *Chaoborus* larvae than the other cladocerans [50]. The presence of roach made the planktonic communities gradually deviate from those of the fishless communities with a shift in the dominant taxon of zooplankton from daphnids to bosminids whose abundance greatly increased. Visual-feeding fish like roach tend to prefer large zooplankton [51] and it might be that selective predation on the two largest taxa: *Chaoborus* larvae and daphnids, has released bosminids from predation and competition. Alteration in the daphnids – bosminids balance is symptomatic of fish presence [52]. To a lesser extent, copepods and Chydoridae have also benefited from the roach-induced decrease in daphnids. This might be explained by the greater availability of food resources like green algae, which slightly increased in the presence of roach, but also rotifers that represent another important taxon of freshwater zooplankton. Due to their small size, we did not count the number of rotifers but they are known to increase in the presence of fish because of the removal of large cladocerans [53]. The slight increase in green algae in the presence of roach is consistent with the trophic cascade hypothesis: fish have a negative direct effect on zooplankton (here daphnids) and indirectly benefit phytoplankton that is released from grazing [34]. Another way roach can influence the planktonic communities is through the modulation of diel migration patterns. Indeed, some taxa migrate to the bottom under chemical cues by predators and become less frequent in the pelagic realm [54].

Motorboat noise did not alter the top-down structuring effect of roach on the planktonic communities and particularly the shift from daphnids to bosminids. This suggests that chronic exposure to noise had no effect on the feeding behaviour of roach, which is consistent with the absence of difference in growth rate between the two noise conditions. This is also consistent with the total number of prey eaten recorded during the predation tests, which was significantly reduced by motorboat noise for the roach that never experienced boat noise before but not for those pre-exposed to boat noise in the mesocosms. Weakening of the response to noise after repeated exposure has been reported in other fish species [38,55,56], and might reflect habituation through associative learning: naïve animals first allocate attention to noise at the expense of other activities like feeding, and then resume normal behaviour as they learn that it is not associated with any threat. However, when looking closer to what has been eaten, we found motorboat noise to elicit selective preference for daphnids over *Chaoborus* larvae even for roach pre-exposed to boat noise in the mesocosm. This persistent response could find its origin in behaviour. Concerning invertebrates, although we did not record their behaviour, we know from past investigations that motorboat noise did not alter the mobility of daphnids [27] but triggers body rotations in *Chaoborus* larvae [38], interpreted as an anti-predatory response [57,58] that could have driven the choice of roach towards daphnids. Concerning roach behaviour, we found no alteration in mobility and no evidence for any avoidance of the sound source, but the area occupied by the three individuals was larger under motorboat noise. This effect seems to be persistent as it was also observed with the roach pre-exposed to boat noise in the mesocosms. Similarly, playback of pile driving was found to make juveniles of seabass les cohesive [59]. Noise could mask the perception of nearest neighbours’ movements through the lateral line or impair the ability to process sensory information as a consequence of stress and/or distraction [59]. Compared to stress or distraction, masking does not weaken with repeated exposure. This could explain why the reduced group cohesion was also observed in the roach that experienced motorboat noise in the mesocosms. Disruption of group cohesion could ultimately compromise the benefits of grouping associated with the dilution and confusion effects [60]. Regarding feeding, we can expect the strength of intra-specific competition to decrease with the distance between individuals. Together with the lesser catchability of moving *Chaoborus* larvae, this could explain why the roach showed selective preference for daphnids under motorboat noise. At the level of communities, a selective preference for daphnids, which were the main grazers, should have strengthened the trophic cascade, what we did not observe. The change in feeding we found in the aquaria may not have been strong enough to be detected in the mesocosms or maybe does not occur in a larger and more complex environment where other prey items are available.

In the fishless mesocosms, adding motorboat noise induced a small but detectable deviation from the control communities. This is interesting but also tricky to interpret since very little is known on the response of freshwater plankton to chronic anthropogenic noise. *Chaoborus* larvae occupied the highest trophic level of the fishless communities and we know that they are sensitive to motorboat sounds with more body rotations [38], interpreted as an anti-predatory response [58]. If noise also interferes with prey processing and reduces the capture efficiency of *Chaoborus* larvae, then it could be beneficial to small zooplankton. Noise could also trigger vertical migration to the bottom, as chemical cues from predators do [54], making some taxa like ostracods less detectable in the pelagic realm. Additional long-term investigations in mesocosms but also *in situ* are needed to better understand the response of planktonic communities to chronic anthropogenic noise.

Our investigation illustrates how extrapolating the impact of anthropogenic noise from individual responses to complex communities if far from obvious. Although we observed persistent alterations in roach behaviour with less group cohesion and altered feeding preference, these effects did not propagate downward along the food chain through trait-mediated cascading effects. A valuable perspective would be to study the dynamics of roach under chronic anthropogenic noise to test whether the behavioural responses we observed ultimately decrease survival and/or reproductive success [61], and result in density-mediated cascading effects.

## ACKNOWLEDGEMENTS

We thank the CEREEP Ecotron Ile-De-France (CNRS/ENS UMS 3194) for access to the PLANAQUA experimental facilities and help during the experiments. A special thanks to Beatriz Decencière and Jacques Mériguet for their help in the realization of the experiment. We are also grateful to Mathieu Mullot for assistance during the PRC analysis.

## ETHIC STATEMENTS

All the procedures were conducted in accordance with appropriate European (*Directive* 2010/63/EU) and French national guidelines, permits, and regulations regarding animal care and experimental use (Approval no. C42-218-0901).

## DATA AVAILABILITY

The data that support the findings of this study are open available in Zenodo at https://doi.org/10.5281/zenodo.6760791.

## REFERENCES

1. Fretwell SD. 1987 Food Chain Dynamics: The Central Theory of Ecology? Oikos 50, 291–301. (doi:10.2307/3565489)

2. Ripple WJ et al. 2016 What is a Trophic Cascade? Trends in Ecology & Evolution 31, 842–849. (doi:10.1016/j.tree.2016.08.010)

3. Schmitz OJ, Krivan V, Ovadia O. 2004 Trophic cascades: the primacy of trait-mediated indirect interactions. Ecology Letters 7, 153–163. (doi:10.1111/j.1461-0248.2003.00560.x)

4. Su H, Feng Y, Chen J, Chen J, Ma S, Fang J, Xie P. 2021 Determinants of trophic cascade strength in freshwater ecosystems: a global analysis. Ecology 102, e03370. (doi:10.1002/ecy.3370)

5. Hebblewhite M. 2005 Predation by Wolves Interacts with the North Pacific Oscillation (NPO) on a Western North American Elk Population. Journal of Animal Ecology 74, 226–233.

6. Cheng BS, Grosholz ED. 2016 Environmental stress mediates trophic cascade strength and resistance to invasion. Ecosphere 7, e01247. (doi:10.1002/ecs2.1247)

7. Duchet C et al. 2018 Pesticide-mediated trophic cascade and an ecological trap for mosquitoes. Ecosphere 9, e02179. (doi:10.1002/ecs2.2179)

8. Sordello R, Ratel O, Flamerie De Lachapelle F, Leger C, Dambry A, Vanpeene S. 2020 Evidence of the impact of noise pollution on biodiversity: a systematic map. Environmental Evidence 9, 20. (doi:10.1186/s13750-020-00202-y)

9. Duarte CM et al. 2021 The soundscape of the Anthropocene ocean. Science 371, eaba4658. (doi:10.1126/science.aba4658)

10. Buxton RT, McKenna MF, Mennitt D, Fristrup K, Crooks K, Angeloni L, Wittemyer G. 2017 Noise pollution is pervasive in U.S. protected areas. Science 356, 531–533. (doi:10.1126/science.aah4783)

11. Kight CR, Swaddle JP. 2011 How and why environmental noise impacts animals: an integrative, mechanistic review. Ecology Letters 14, 1052–1061. (doi:10.1111/j.1461-0248.2011.01664.x)

12. Francis CD, Barber JR. 2013 A framework for understanding noise impacts on wildlife: an urgent conservation priority. Frontiers in Ecology and the Environment 11, 305–313. (doi:10.1890/120183)

13. Shannon G et al. 2016 A synthesis of two decades of research documenting the effects of noise on wildlife. Biological Reviews 91, 982–1005. (doi:10.1111/brv.12207)

14. Francis CD, Kleist NJ, Ortega CP, Cruz A. 2012 Noise pollution alters ecological services: enhanced pollination and disrupted seed dispersal. Proceedings of the Royal Society B: Biological Sciences 279, 2727–2735. (doi:10.1098/rspb.2012.0230)

15. Phillips JN, Termondt SE, Francis CD. 2021 Long-term noise pollution affects seedling recruitment and community composition, with negative effects persisting after removal. Proceedings of the Royal Society B: Biological Sciences 288, 20202906. (doi:10.1098/rspb.2020.2906)

16. Barton BT, Hodge ME, Speights CJ, Autrey AM, Lashley MA, Klink VP. 2018 Testing the AC/DC hypothesis: Rock and roll is noise pollution and weakens a trophic cascade. Ecology and Evolution 8, 7649–7656. (doi:10.1002/ece3.4273)

17. Dudgeon D et al. 2006 Freshwater biodiversity: importance, threats, status and conservation challenges. Biological Reviews 81, 163–182. (doi:10.1017/S1464793105006950)

18. Dudgeon D. 2019 Multiple threats imperil freshwater biodiversity in the Anthropocene. Current Biology 29, R960–R967. (doi:10.1016/j.cub.2019.08.002)

19. Maasri A et al. 2022 A global agenda for advancing freshwater biodiversity research. Ecology Letters 25, 255–263. (doi:10.1111/ele.13931)

20. Slabbekoorn H, Bouton N, van Opzeeland I, Coers A, ten Cate C, Popper AN. 2010 A noisy spring: the impact of globally rising underwater sound levels on fish. Trends in Ecology & Evolution 25, 419–427. (doi:10.1016/j.tree.2010.04.005)

21. Kunc HP, McLaughlin KE, Schmidt R. 2016 Aquatic noise pollution: implications for individuals, populations, and ecosystems. Proceedings of the Royal Society B: Biological Sciences 283, 20160839. (doi:10.1098/rspb.2016.0839)

22. Cox K, Brennan LP, Gerwing TG, Dudas SE, Juanes F. 2018 Sound the alarm: A meta-analysis on the effect of aquatic noise on fish behavior and physiology. Global Change Biology 24, 3105–3116. (doi:10.1111/gcb.14106)

23. Mickle MF, Higgs DM. 2018 Integrating techniques: a review of the effects of anthropogenic noise on freshwater fish. Canadian Journal of Fisheries and Aquatic Sciences 75, 1534–1541. (doi:10.1139/cjfas-2017-0245)

24. Lampert W. 2006 Daphnia: model herbivore, predator and prey. Polish Journal of Ecology 54, 607–620.

25. Reynolds CS. 2011 Daphnia: Development of Model Organism in Ecology and Evolution - 2011. Freshwater Reviews 4, 85–87. (doi:10.1608/FRJ-4.1.425)

26. Tidau S, Briffa M. 2016 Review on behavioral impacts of aquatic noise on crustaceans. Proceedings of Meetings on Acoustics 27, 010028. (doi:10.1121/2.0000302)

27. Sabet SS, Neo YY, Slabbekoorn H. 2015 The effect of temporal variation in sound exposure on swimming and foraging behaviour of captive zebrafish. Animal Behaviour 107, 49–60. (doi:10.1016/j.anbehav.2015.05.022)

28. Sabet SS, Karnagh SA, Azbari FZ. 2019 Experimental test of sound and light exposure on water flea swimming behaviour. Proceedings of Meetings on Acoustics 37, 010015. (doi:10.1121/2.0001270)

29. Hawkins A, Myrberg A. 1983 Hearing and sound communication underwater. Bioacoustics, a comparative approach, Academic Press, London, 347–405.

30. Popper AN, Fay RR. 2011 Rethinking sound detection by fishes. Hearing Research 273, 25–36. (doi:10.1016/j.heares.2009.12.023)

31. Andersson MH, Dock-Akerman E, Ubral-Hedenberg R, Ohman MC, Sigray P. 2007 Swimming behavior of roach (Rutilus rutilus) and three-spined stickleback (Gasterosteus aculeatus) in response to wind power noise and single-tone frequencies. Ambio 36, 636–638. (doi:10.1579/0044-7447(2007)36[636:sborrr]2.0.co;2)

32. Amoser S, Wysocki LE, Ladich F. 2004 Noise emission during the first powerboat race in an Alpine lake and potential impact on fish communities. The Journal of the Acoustical Society of America 116, 3789–3797. (doi:10.1121/1.1808219)

33. Magnhagen C, Johansson K, Sigray P. 2017 Effects of motorboat noise on foraging behaviour in Eurasian perch and roach: a field experiment. Marine Ecology Progress Series 564, 115–125. (doi:10.3354/meps11997)

34. Bertolo A, Lacroix G, Lescher-Moutouš F, Sala S. 1999 Effects of physical refuges on fish–plankton interactions. Freshwater Biology 41, 795–808. (doi:10.1046/j.1365-2427.1999.00424.x)

35. Haney JF, et al. 2013 ‘An-Image-based Key to the Zooplankton of North America’ version 5.0 released in 2013. University of New Hampshire Center for Freshwater Biology. See http://cfb.unh.edu/cfbkey/html/index.html.

36. Winterbourn MJ, Winterbourn G, Katharine L, Katharine D, Heath C. 1989 Guide to the aquatic insects of New Zealand. Entomological Society of New Zealand Auckland.

37. Spitze K. 1991 Chaoborus predation and life-history evolution in Daphnia pulex: temporal pattern of population diversity, fitness, and mean life history. Evolution 45, 82–92. (doi:10.1111/j.1558-5646.1991.tb05268.x)

38. Rojas E, Thévenin S, Montes G, Boyer N, Médoc V. 2021 From distraction to habituation: Ecological and behavioural responses of invasive fish to anthropogenic noise. Freshwater Biology 66, 1606–1618. (doi:10.1111/fwb.13778)

39. Sueur J, Aubin T, Simonis C. 2008 Seewave, a free modular tool for sound analysis and synthesis. Bioacoustics 18, 213–226. (doi:10.1080/09524622.2008.9753600)

40. R Core Team. 2021 R: A Language and Environment for Statistical Computing. Vienna, Austria: R Foundation for Statistical Computing. See https://www.R-project.org/.

41. Oksanen J et al. 2013 vegan: Community Ecology Package. Community ecology package, version 2, 1–295.

42. Van den Brink PJ, Braak CJFT. 1999 Principal response curves: Analysis of time-dependent multivariate responses of biological community to stress. Environmental Toxicology and Chemistry 18, 138–148. (doi:10.1002/etc.5620180207)

43. Legendre P, Gallagher ED. 2001 Ecologically meaningful transformations for ordination of species data. Oecologia 129, 271–280. (doi:10.1007/s004420100716)

44. Bates D, Maechler M, Bolker B, Walker S. 2015 Fitting Linear Mixed-Effects Models Using lme4. Journal of Statistical Software 67, 1–48. (doi:10.18637/jss.v067.i01)

45. Manly BFJ. 1974 A Model for Certain Types of Selection Experiments. Biometrics 30, 281–294. (doi:10.2307/2529649)

46. Chesson J. 1983 The Estimation and Analysis of Preference and Its Relatioship to Foraging Models. Ecology 64, 1297–1304. (doi:10.2307/1937838)

47. Manly BFJ. 1995 A Note on the Analysis of Species Co-Occurrences. Ecology 76, 1109–1115. (doi:10.2307/1940919)

48. Barton K. 2009 MuMIn: Multi-Model Inference. See http://r-forge.r-project.org/projects/mumin/.

49. Elser MM, Ende CN von, Sorrano P, Carpenter SR. 1987 Chaoborus populations: response to food web manipulation and potential effects on zooplankton communities. Canadian Journal of Zoology 65, 2846–2852. (doi:10.1139/z87-433)

50. Jäger IS, Hölker F, Flöder S, Walz N. 2011 Impact of Chaoborus flavicans -Predation on the Zooplankton in a Mesotrophic Lake – a Three Year Study. International Review of Hydrobiology 96, 191–208. (doi:10.1002/iroh.201011253)

51. Jarolím O, Kubečka J, Čech M, Vašek M, Peterka J, Matěna J. 2010 Sinusoidal swimming in fishes: the role of season, density of large zooplankton, fish length, time of the day, weather condition and solar radiation. Hydrobiologia 654, 253–265. (doi:10.1007/s10750-010-0398-1)

52. Post JR, McQueen DJ. 1987 The impact of planktivorous flsh on the structure of a plankton community. Freshwater Biology 17, 79–89. (doi:10.1111/j.1365-2427.1987.tb01030.x)

53. Gilbert JJ. 1988 Suppression of rotifer populations by Daphnia: A review of the evidence, the mechanisms, and the effects on zooplankton community structure1. Limnology and Oceanography 33, 1286–1303. (doi:10.4319/lo.1988.33.6.1286)

54. Cohen JH, Forward RB. 2016 Zooplankton diel vertical migration - A review of proximate control. Oceanography and marine biology 47, 89–122.

55. Nedelec SL, Mills SC, Lecchini D, Nedelec B, Simpson SD, Radford AN. 2016 Repeated exposure to noise increases tolerance in a coral reef fish. Environmental Pollution 216, 428–436. (doi:10.1016/j.envpol.2016.05.058)

56. Neo YY, Hubert J, Bolle LJ, Winter HV, Slabbekoorn H. 2018 European seabass respond more strongly to noise exposure at night and habituate over repeated trials of sound exposure. Environmental Pollution 239, 367–374. (doi:10.1016/j.envpol.2018.04.018)

57. Berendonk TU, O’Brien WJ. 1996 Movement response of Chaoborus to chemicals from a predator and prey. Limnology and Oceanography 41, 1829–1832. (doi:10.4319/lo.1996.41.8.1829)

58. Burrows M, Dorosenko M. 2014 Rapid swimming and escape movements in the aquatic larvae and pupae of the phantom midge Chaoborus crystallinus. Journal of Experimental Biology 217, 2468–2479. (doi:10.1242/jeb.102483)

59. Herbert-Read JE, Kremer L, Bruintjes R, Radford AN, Ioannou CC. 2017 Anthropogenic noise pollution from pile-driving disrupts the structure and dynamics of fish shoals. Proceedings of the Royal Society B: Biological Sciences 284, 20171627. (doi:10.1098/rspb.2017.1627)

60. Krause J, Ruxton GD, Ruxton G, Ruxton IG. 2002 Living in groups. Oxford University Press.

61. Amorim MCP et al. 2022 Boat noise impacts Lusitanian toadfish breeding males and reproductive outcome. Science of The Total Environment 830, 154735. (doi:10.1016/j.scitotenv.2022.154735)

